# Dynamic Network Analysis of the 4D Nucleome

**DOI:** 10.1101/268318

**Authors:** Sijia Liu, Pin-Yu Chen, Alfred Hero, Indika Rajapakse

## Abstract

**Motivation:** For many biological systems, it is essential to capture simultaneously the function, structure, and dynamics in order to form a comprehensive understanding of underlying phenomena. The dynamical interaction between 3D genome spatial structure and transcriptional activity creates a genomic signature that we refer to as the four-dimensional organization of the nucleus, or 4D Nucleome (4DN). The study of 4DN requires assessment of genome-wide structure and gene expression as well as development of new approaches for data analysis.

**Results:** We propose a dynamic multilayer network approach to study the co-evolution of form and function in the 4D Nucleome. We model the dynamic biological system as a temporal network with node dynamics, where the network topology is captured by chromosome conformation (Hi-C), and the function of a node is measured by RNA sequencing (RNA-seq). Network-based approaches such as von Neumann graph entropy, network centrality, and multilayer network theory are applied to reveal universal patterns of the dynamic genome. Our model integrates knowledge of genome structure and gene expression along with temporal evolution and leads to a description of genome behavior on a system wide level. We illustrate the benefits of our model via a real biological dataset on MYOD1-mediated reprogramming of human fibroblasts into the myogenic lineage. We show that our methods enable better predictions on form-function relationships and refine our understanding on how cell dynamics change during cellular reprogramming.

**Availability:** The software is available upon request.

**Supplementary information:** See Supplementary Material.

## 1 Introduction

The 4D Nucleome (4DN) reflects a dynamic interplay between 3D genome architecture (form) and transcription (function), as well as its impact on phenotype (Dekker *et al.*, 2017; Dixon *et al.*, 2015; Fortin and Hansen, 2015; Chen *et al.*, 2015; Krijger *et al.*, 2016). Genome technologies such as genome-wide chromosome conformation capture (Hi-C) (Lieberman-Aiden *et al.*, 2009) are yielding ever higher resolution data that give a more complete picture of the 4DN, allowing us to refine the roles of cell types, lineage differentiation, and pathological contributions of cells in different diseases. With the aid of Hi-C and RNA-sequencing (RNA-seq) data captured over time, we propose a mathematical foundation for the study of 4DN. Specifically, we introduce a dynamic multilayer network approach to study the co-evolution of form and function in the 4DN. We view the genome as a dynamic network, where at each time, the network topology (chromosomal contacts) and function of nodes (genes) are determined by Hi-C contact map and gene expression, respectively.

Some recent research efforts have been made to explore the form-function relationship (Lieberman-Aiden *et al.*, 2009; Dixon *et al.*, 2015, 2012; Chen *et al.*, 2015). One important finding was that Hi-C contact maps can be used to divide the genome into two major compartments, termed A and B (Lieberman-Aiden *et al.*, 2009), where compartment A is associated with open chromatin (transcriptionally active), and compartment B is associated with closed chromatin (transcriptionally inactive). Interestingly, the work (Chen *et al.*, 2015) showed that A/B compartments can be interpreted as distinct connected components of the network representation of a genome. It was also shown in (Lieberman-Aiden *et al.*, 2009; Dixon *et al.*, 2015, 2012) that the structurally defined regions (i.e. A/B compartments) change over time to facilitate different expression patterns of a new cell state. The previous findings suggest that there exists a certain relationship between genomic form and function, which can be furthermapped to the relationship between the network topology and the function of the nodes in the network.

Different from the existing literature (Lieberman-Aiden *et al.*, 2009; Dekker *et al.*, 2013; Dixon *et al.*, 2015; Fortin and Hansen, 2015; Chen *et al.*, 2015; Krijger *et al.*, 2016), the dynamic network point of view allows us to extract multiple topological properties of the genome over time, and facilitates quantitative integration with gene expression data. In this paper, network-based approaches including von Neumann graph entropy, network centrality, and multilayer network theory are adopted to study the genome dynamics. We show that the network entropy provides an efficient way to track the amount of uncertainty of the genome as it evolves over time, and the use of network centrality helps to build biological connections between the genome structure and network properties. With the aid of multilayer network theory, we can further reveal details on nuclear reorganization over time, e.g., how contacts between genes evolve during cell development.

Our proposed dynamic multilayer network approach is applied to a real biological dataset on MYOD1-mediated fibroblast-to-muscle reprogramming (Liu *et al.*, 2017). Compared to a normal cell development process, e.g., fibroblast proliferation (Chen *et al.*, 2015), cellular reprogramming is a process of converting one cell (e.g., fibroblast) to another (e.g., muscle) through overexpression of certain transcription factors (TFs; e.g., MYOD1). Although cellular reprogramming has been studied experimentally (Weintraub *et al.*, 1989; Weintraub, 1993; Takahashi *et al.*, 2007; Ronquist *et al.*, 2017; Liu *et al.*, 2017), the question of how genome architecture and transcription co-evolve as a cell adopts a new identity is not well understood. Understanding such system-level genome dynamics may allow more sophisticated reprogramming strategies that could be used in cell therapeutics.

Our dynamic multilayer network approach to analyzing the 4DN results in two main biological findings that are consistent with results reported in (Liu *et al.*, 2017). First, we confirm that cellular reprogramming undergoes a critical transition point, given by an intermediate highly entropic state. This state can be represented as a bifurcation point in a dynamical system (Liu *et al.*, 2017), where one stable equilibrium (fibroblast) bifurcates to two stable equilibria (fibroblast and muscle). Second, we find that the chromatin reorganization significantly changes *before* transcriptional changes occur during the reprogramming process. In other words, form precedes function. Moreover, we uncover small but influential set of genes that governs cell cycle dynamics. Our proposed method will lead to improved understanding of form-function dynamics, which is key to advancing cellular reprogramming strategies. This perspective can have a broad translational impact spanning cancer cell biology and regenerative medicine. Although our experimental study is restricted to MYOD1-mediated reprogramming, the proposed methods are applicable to form-function analysis of any dynamic biological system.

## 2 Preliminaries: Genomic Form and Function

In this section, we provide background on Hi-C and RNA-seq data to capture genomic form and function.

The Hi-C technique was first introduced by (Lieberman-Aiden *et al.*, 2009) to represent the genome organization through a 2D contact map. Specifically, Hi-C evaluates long-range interactions between pairs of segments delimited by specific cutting sites using spatially constrained ligation. This leads to a fragment level contact map, in terms of a fragment read table. Each row of this table indicates a ligated pair of fragments from the genome, with the coordinates of both fragments. Then, given any two regions in the genome, the contacts of these two regions can be identified by searching over the fragment read table and counting the number of pairs that are located within these regions. The Hi-C contact maps are often generated by binning fragments into fixed resolution bins such as 1 megabase (Mb) and 100 kilobase (Kb) bins. Both of 1 Mb and 100 Kb binned Hi-C contact maps will be used in this paper.

Mathematically, a Hi-C contact map can be expressed as a square nonnegative symmetric matrix **H**, where the (*i, j*)-th entry *H_ij_* denotes the number of contacts between loci *i* and *j* at a given scale. Since raw Hi-C matrices suffer from biases due to the effect of 1D genome mapping: closely spaced loci are likely to have large Hi-C read counts regardless of their actual 3D conformation. Therefore, a raw Hi-C measurement is diagonally dominant and tapered. To mitigate this bias, we normalize Hi-C matrices using a Toeplitz normalization method (Chen *et al.*, 2016) prior to data analysis.

In addition to Hi-C data, RNA-seq measures the expression level of each gene in the unit of Fragments Per Kilobase of transcript per Million mapped reads (FPKM). As an example of dynamic biological systems, we consider a cellular reprogramming experiment (Liu *et al.*, 2017), which converts primary human fibroblasts into the myogenic lineage using the master regulator MYOD1. Fig. 1 provides an illustration of the experiment and the result of analysis, the graph at the right, that is the subject of this paper. A skin cell is reprogrammed to a skeletal muscle cell over a 56-hour (hr) period, 8 time samples (0, 8,…, 56 hrs) are taken to capture the time-evolving genome architecture and transcription via Hi-C and RNA-seq. The Hi-C library, RNA-seq library, data collection, and raw data processing were all performed by our laboratory; see (Liu *et al.*, 2017, Methods) for more details.

**Fig. 1.**
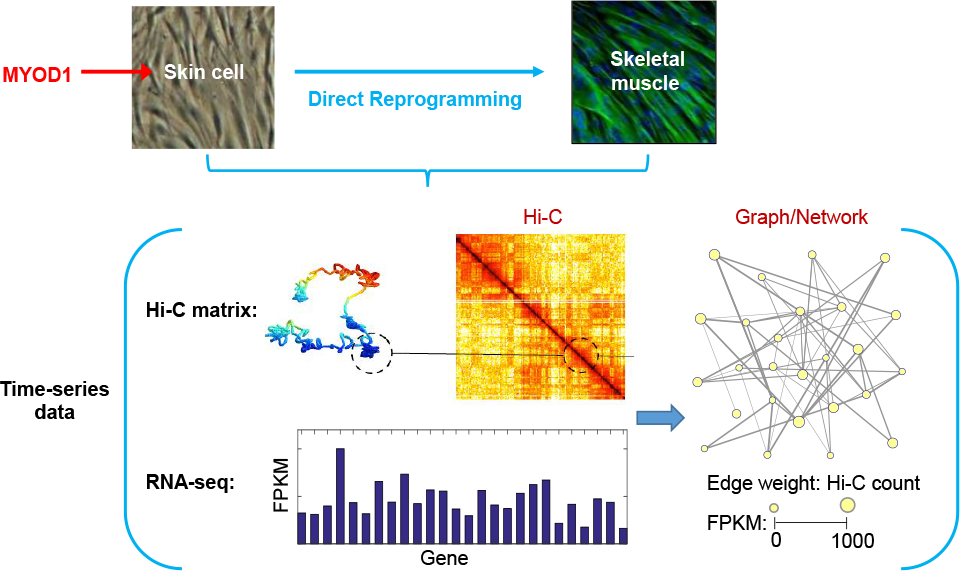
Illustration of the cellular reprogramming experiment used in this paper to
demonstrate the proposed integrated Hi-C and RNA-seq graph extraction algorithm.

## 3 Dynamic Multilayer Network Analysis for 4DN

In this section, we develop the proposed network-based approach to integrate the dynamic genome architecture (form) and transcription (function) from experimental data.

### 3.1 Network representation

The availability of Hi-C data allows us to study the spatial organization of a genome from a network (or graph) point of view (Fig. 1). Specifically, nodes of the network correspond to genomic loci that can be partitioned at different scales such as bin or gene resolution. Edges of the network correspond to pairs of loci that are in physical contact, with edge weights given by values in the Hi-C matrix. Gene expression is quantified by RNA-seq values, which can be interpreted as attributes of nodes in the network. Mathematically, let 𝒢=(𝒱, *ε*, **W**; y) denote the network representation of genomic form and function, where 𝒱 is the node set (genomic loci) with cardinality |𝒱|=*n*, *ε* is the edge set, **W** ∈ ℝ^*n*×*n*^ is a weighted adjacency matrix with *W_ij_*=*H_ij_* if *i*≠*j* and *W_ii_*=0 by convention, **y** ∈ ℝ^*n*^ is the vector of gene expression, and ℝ denotes the set of real numbers. Using the above notation, adynamic network over a time horizon of length T can be described by a graph sequence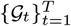

The quantitative study of 𝒢 is often performed over its graph Laplacian matrix **L** = **D** − **W**, where **D** = diag(**W1**) is the degree matrix of G with diagonal vector **W1**, and **1** is the vector of all ones. The graph Laplacian matrix **L** is positive semidefinite and its normalized version **L**_norm_ = **I** − **D**^−1/2^**WD**^−1/2^ is called the normalized graph Laplacian matrix. From spectral graph theory (Chung, 1997), the second smallest eigenvalue of **L** (or **L**_norm_), known as the Fiedler number (FN), characterizes the network connectivity. Note that G is connected (namely, there exists a path between every pair of distinct nodes) if and only if the FN is nonzero. The eigenvector associated with the FN is called the Fiedler vector, which encodes the community information of a network (Chen *et al.*, 2016).

For the integrative analysis of genomic form (**W**) and function (y), we encounter two main issues: how to track the dynamical state of a dynamic biological system and how to efficiently and quantitatively integrate form-function data. In what follows, we propose possible solutions to address these problems.

### 3.2 Von Neumann Graph Entropy

Von Neumann graph entropy (VNGE) is the von Neumann entropy of a graph and it allows us to quantify the amount of information encoded in the genome architecture, and to evaluate differences or similarities between dynamic biological networks (Braunstein *et al.*, 2006; Passerini and Severini, 2008; Han *et al.*, 2012). Note that von Neumann entropy was originally used to measure the incompressible information content of a quantum source, which can characterize the departure of a dynamical system from a pure state with zero entropy (Anand *et al.*, 2011). This provides the motivation for using VNGE to measure the complexity of a dynamic biological system.

The VNGE (*V*) of a network 𝒢 is determined by the spectrum of a scaled graph Laplacian matrix **L**_*c*_ : = *c***L** with *c* = 1 / trace(**L**),

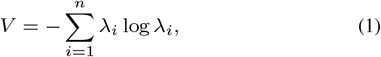

where trace(․) denotes the trace operator of a matrix, and 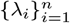 are eigenvalues of **L**_*c*_. It is clear from (1) that VNGE can be interpreted as the Shannon entropy of the probability distribution represented by 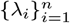under the facts that λ_*i*_ ≥ 0 for any *i* and 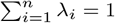.

With the aid of VNGE, we are able to track the amount of uncertainty of a dynamic genome. The network entropy can also be linked with controllability gramian of a biological system (Rajapakse *et al.*, 2012). A highly entropic state of the system suggests a high degree of controllability, or receptivity to external influence. In (1), VNGE requires the full eigenspectrum of the graph Laplacian matrix, and thus has cubic computational complexity *O*(*n*^3^). In order to reduce this complexity, we propose an approximate VNGE that scales better with the network size *n*.

Considering a quadratic approximation *Q* of *V* based on the Taylor series expansion of 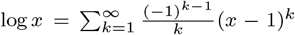 at *x* = 1, we obtain

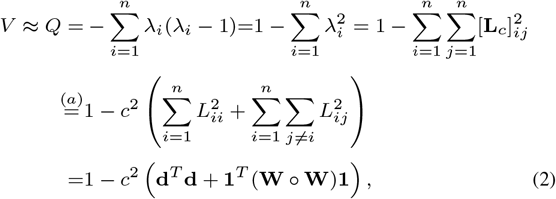

where the equality (*a*) holds due to the definition of **L**_*c*_, [**A**]*_ij_* (or *A*_*ij*_) represents the (*i, j*)-th entry of a matrix **A**, **d** is the diagonal vector of **L**, **W** is the weighted adjacency matrix, and ○ denotes the entrywise (Hadamard) product. Compared to (1), the quadratic approximation (2) yields an improved complexity *O*(*n*^2^). Moreover, under mild conditions, the asymptotic consistency of *Q* with respect to *V* is guaranteed: *Q* log *n* → *V* as *n* → ∞ (see details in Supplementary Material). In Sec. 4, we will show that VNGE helps to identify a bifurcation event that delineates emergence of a new cell identity.

### 3.3 Network Centrality Analysis

With the aid of network centrality, we are able to evaluate the topological influence of genomic loci on the genome. By extracting multiple centrality features from a dynamic network, we can find intrinsic low-dimensional manifolds embedded in form-function data. A number of centrality measures are used, such as degree, eigenvector, clustering coefficient, closeness, betweenness, and hop walk statistics differing in the types of influence that one wishes to emphasize (Newman, 2010). Here we highlight three types of network centrality and discuss their biological interpretations.

**Fig. 1:**
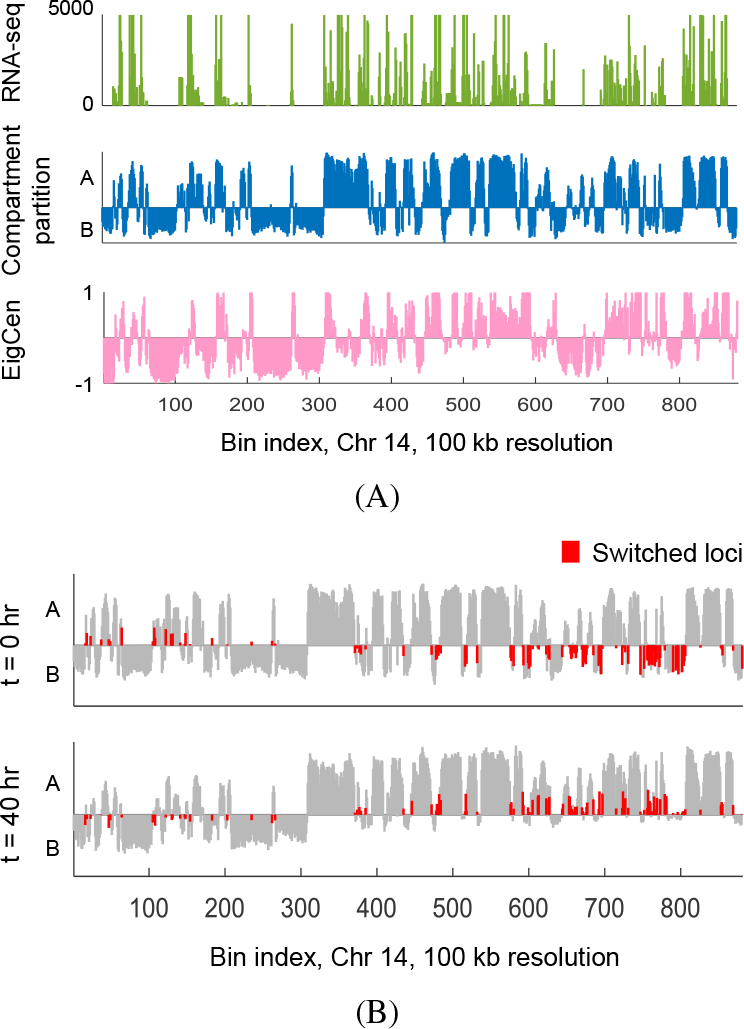
Examples of network centrality and their biological insights. (A) Chromatin compartment partition for chromosome 14. (B) Loci of compartment change (i.e., A/B switch) at 0 and 40 hrs.

*Degree* (*Deg*) of node *i* is defined as the sum of edge weights (namely, Hi-C contacts) associated with it, Deg(*i*) = [**W1**]_*i*_ = *L*_*ii*_, where we recall that **W** and **L** are graph adjacency and Laplacian matrices, respectively. Deg characterizes the local connectedness of a node (namely, a genomic locus) as measured by interactions with other nodes. From a biological perspective a node with high degree centrality is a locus in contact with many other loci on the genome (form similarity) or a gene whose expression profile is very similar to the profiles of many other genes(functional similarity).

*Eigenvector centrality* (*EigCen*) is defined as the eigenvector of the adjacency matrix **W** associated with its largest positive eigenvalue λ_max_. The eigenvector centrality of node *i* is given by EigCen(*i*) = [**v**]_*i*_ satisfying λ_max_**v** = **Wv**. EigCen is a neighborhood connectedness property in which a node has high centrality if many of its neighbors also have high centrality. From a biological perspective, EigCen identifies structurally defined regions of active/inactive gene expression, since it encodes clustering in a network (Ng *et al.*, 2001); see Fig.2A for an example. It has been shown in (Liu *et al.*, 2017) that EigCen yields higher correlation with transcriptional activity than chromatin compartment partition (i.e., A/B compartments), which is often defined using the first principal component of spatial correlation matrix of Hi-C contact map (Lieberman-Aiden *et al.*, 2009).

*Betweenness* (*Betw*) measures the fraction of shortest paths passing through a node relative to the total number of shortest paths in the network (Freeman, 1977). The betweenness of node *i* is defined as 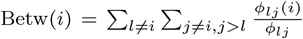 where *Φ*_*lj*_ the total number of shortest paths from node *l* to *j*, and *Φ*_*lj*_ is the number of such shortest paths passing through node *i*. Compared to Deg and EigCen, Betw is a global connectedness measure since it quantifies the number of times a node acts as a bridge along the shortest path between two other nodes. From a biological perspective, Betw recognizes the switched regions (Fig.2B) of chromatin compartments over time in the sense that Betw becomes significantly higher than other centrality measures during these switching transitions. (Liu *et al.*, 2017). That is because many switches (more than 70%) appear at boundaries of chromatin compartments, which serve as bridges between A/B compartments and thus switches become bottleneck nodes connecting one compartment to another.

The biological interpretation of network centrality motivates the extraction of multiple topological properties of the genome using Hi-C data. The extracted feature vectors can then be combined with gene expression, leading to a form-function feature matrix **X**_*t*_ ∈ ℝ^*n*×*p*^ at time *t*, where *p* is the number of form-function features. We can employ dimensionality reduction techniques (Van Der Maaten *et al.*, 2009) to find the intrinsic low-dimensional manifold embedded in 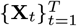. This leads to a low-dimensional data representation **Y**_*t*_ ∈ℝ^*n*×*l*^ with *l* the feature embedding dimension, *l*≤*p*. Each column of the matrix Y_*t*_ is an observed instance of the dimension reduced data sample, called the *state*, at time *t*.

In order to better track the state of a dynamic genome, we fit the data **Y**_*t*_ to a minimum volume ellipsoid (MVE), representing a certain confidence region containing the state. The MVE estimate at time *t* is acquired by solving the convex program

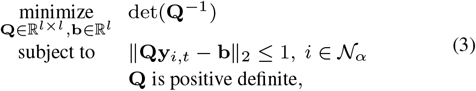

where **Q** and **b** are optimization variables, 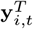 is the *i*th rowvector of **Y**_*t*_,and 𝒩_*α*_ denotes the set of data within *α* confidence region, determined by Mahalanobis distances of data below *α* = 97.5% quantile of the chisquare distribution with *l* degrees of freedom (Van Aelst and Rousseeuw,2009). The rationale behind problem (3) is that the matrix **P** := **Q**^2^ andthe vector **c** := **Q**^−1^**b** define the ellipsoid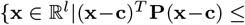1} where the determinant of **P** is inversely proportional to the ellipsoid volume (Boyd and Vandenberghe, 2004). Using a dimension reduction approach we efficiently solve (3) using semidefinite programming (Boyd and Vandenberghe, 2004) under a reduced feature space with small *l*.

Based on (3), we can further adopt Kullback-Leibler (KL) divergence(Kullback and Leibler, 1951) to measure the information divergence of different cell states. Let 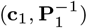 and 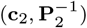 be two MVEs of the state of the form-function data, where we recall that 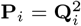 and 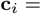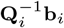 for *i* = 1,2. The symmetrized KL divergence between the states can be defined as SKL_1,2_ = (KL_1,2_ + KL_2,1_)/2, where KL_*i*,*j*_ is the KL divergence

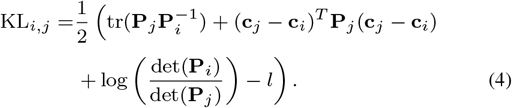

The symmetrized KL divergence (4) is a measure of similarity between the (assumed Gaussian) distributions of the two states. In Sec. 4, we will show that KL helps to detect the critical transition time that marks the change of cell identities.

### 3.4 Multilayer Network Analysis

The dynamic biological network is a multilayer network, where the layers correspond to the snapshots of the network at different time instants (Fig. S1in Supplementary Material). The mulilayer network representation captures the the temporal evolution of the genome.

Recall that nodal degree is an important network centrality measure. However, in a multilayer network, one needs to know not only the degree within a single layer but also how the degree is distributed across different layers. Let 𝒢_*t*_ denote the network of *n* nodes at layer (time) *t*. The degree of node *i* at layer *t* is given by 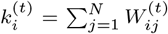 where 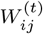 is the (*i, j*)-th entry of the adjacency matrix associated with 𝒢_*t*_. The degree of node *i* for a set of *T* layers is a vector quantity

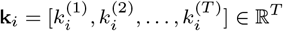

The *overlapping degree* of node *i* across all layers is defined as (Battiston *et al.*, 2014)

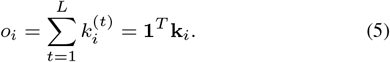

The overlapping degree (5) can be used to identify hubs, namely, nodes with high degree in the network. However, a node that is a hub in one layer may only have few interactions in another layer. Thus, in addition to the overlapping degree, the *multiplex participation coefficient* defines the variation of hub degree across the different layers of the network (Guimera and Amaral, 2005; Battiston *et al.*, 2014),

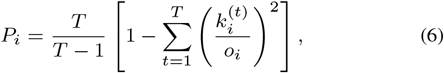

where *P*_*i*_ ∈ [0,1] measures the extent to which the degree of node *i* is uniformly distributed among the *T* layers. If *P*_i_ = 1, then node *i* has exactly the same degree on each layer, namely,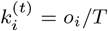.If *P*_*i*_ = 0,all the edges of node *i* are concentrated in just one layer. We will use the multiplex participation coefficient to evaluate the heterogeneity of nodal degrees in dynamic biological networks.

## 4 Experimental Results

In this section, we demonstrate the effectiveness of the proposed dynamic multilayer network analysis of form-function MYOD1-mediated fibroblast-to-muscle reprogramming (Liu *et al.*, 2017). In addition to cellular reprogramming, we also apply our analysis approach to a control study, a fibroblast proliferation dataset (Chen *et al.*, 2015), undergoing normal fibroblast development without the MYOD1 treatment.

### 4.1 Bifurcation: Cell identity transition

First, we useVNGE (1) to assess uncertainty in cellular reprogramming. In Fig. 3A, we present VNGE and its quadratic approximatio (2) of the Hi-C contact maps at 100 kb resolution. The entropic measure is shown under both reprogramming and normal proliferation settings, and is normalized over time in terms of its z-score. First, we note that the approximate VNGE is virtually identical to the VNGE. Second, we observe that the entropic pattern of cellular reprogramming is different from fibroblast proliferation. As we can see from Fig. 3A, the reprogramming process has low entropy at initial and later time points (0 and 56 hrs), demonstrating that the reprogramming system contains two stable basins that encompass a fibroblast state and a myogenic state. The maximum entropy of cellular reprogramming is achieved at 32 hrs. This implies that the reprogramming system undergoes an intermediate highly entropic state, which is in a high degree of receptivity to the external influence of the MYOD1 treatment and induces the cell identity transition.

**Table 1.**
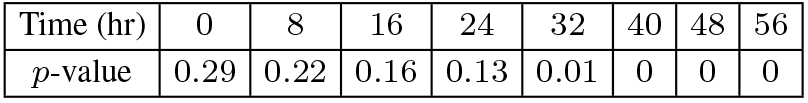
*p*-value for difference between reprogramming and proliferation.

We next show how network centrality can capture structural features of Hi-C contact maps in combination with gene expression, providing an integrative form-function analysis of the 4DN. We apply Laplacian eigenmaps (Supplemental Material), a nonlinear dimensionality reduction technique, to find the low-dimensional representation (the state) of reprogramming and proliferation data. The resulting data configuration is fitted by MVE (3) at each time. The symmetrized version of KL divergence (4) is applied to evaluate the difference between cellular reprogramming and fibroblast proliferation.

In Fig. 4A, we show the timecourse of the symmetrized KL divergence between MVEs of reprogramming and proliferation data. As we can see, there exists a marked divergence at 32 hrs, implying a cell state transition that marks an abrupt shift in the genomic system from its prior state (fibroblast) to a new state (myogenic). This finding is consistent with the entropy results shown in Fig. 3. It was shown in (Liu *et al.*, 2017, Fig.2B) that the transition time (32 hrs) is a bifurcation point in the sense that two simultaneous trends starting from the same state (fibroblast) become separated from each other, toward two stable equilibria (fibroblast and myogenic). Furthermore, in Table 1, we evaluate the significance of the difference between cellular reprogramming and fibroblast proliferation. Here the *p*-value is defined from the Hotelling’s T-squared test associated with the null hypothesis that the centroids of MVEs from reprogramming and proliferation data are identical at a given time point. The detected bifurcation point is further verified experimentally by the timing of the activation of MYOD1 and MYOG in Fig.4B, where MYOD1 and MYOG are key myogenic regulatory factors that initiate the myogenic differentiation.

**Fig. 3.**
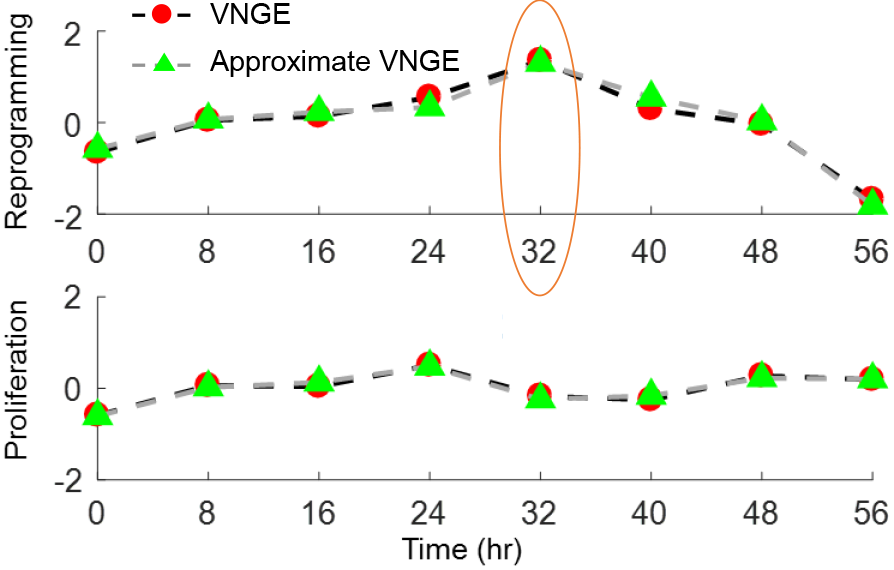
Graph entropy of chromatin organization for cellular reprogramming

**Fig. 4.**
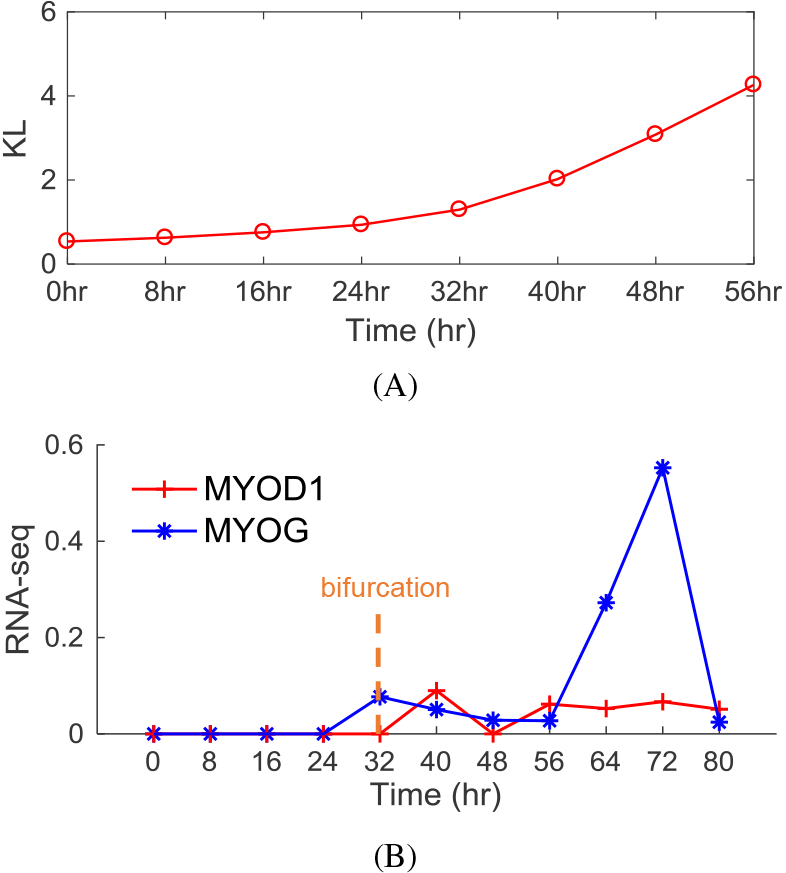
Centrality analysis for bifurcation detection. (A) KL divergence between cellular reprogramming and fibroblast proliferation. (B) Expression ofMYOD1 and MYOG (log scale).

### 4.2 Form precedes function

Given the cell state trajectory, it was unclear whether MYOD1-mediated reprogramming induced rewiring of genome architecture prior mediating muscle gene transcription, or vice versa (Kosak and Groudine, 2004; Rajapakse and Groudine, 2011). To answer this question, we focus on form-function dynamics of genes extracted from Gene Ontology (GO) under three gene modules, fibroblast, muscle and cell cycle (Supplementary Material). Based on genes of interest, we constructed dynamic inter-gene contact maps using fragment-level Hi-C data.

In Fig. 6, we show the temporal change of centrality-based structural features and gene expression at two successive time points. We observe that the largest form change occurs from 8 to 16 hrs while a function change does not occur until at least 32 hrs. This suggests that nuclear reorganization occurs prior to the change of transcriptional activity, that is, form preceded function. In terms of cellular reprogramming, this would imply that chromatin architectural changes facilitate the orchestrated activation of transcriptional networks associated with the adoption of a new cell identity. Furthermore, in Fig. 5 we show a set of 54 cell cycle related genes (Supplementary Table 1) with changes in chromatin structure that proceeded changes in transcriptional activity over the reprogramming time course. Among these genes, 29 maintain increased expression levels identified by edgeR (Robinson *et al.*, 2010) over the time course. From GO annotation (Huang *et al.*, 2008) of the 29 genes, we find 7 genes that are functionally related to the GO term cell cycle arrest, including *ABL1*, *ATM*, *AZK*, *CDKN2A*, *DHCR24*, *GAS1*, and *MAP2K6*. Particularly, the gene *ATM* is one of the master controllers in cell cycle check point signaling pathways (Cortez *et al.*, 2001), and has been shown to be related to DNA stability in skeletal muscle when exposed to laser simulation (Guedes de Almeida *et al.*,2017); others are cell cycle G1 phase controllers (*CDKN2A* and *CHEK2*), or S phase entry blocker (*GAS1*). These results suggest that MYOD1 reorganizes chromatin structure in genomic regions harboring genes involved in cell cycle, and that the increased expression of genes for cell cycle arrest might be required for the cells to exit from the cell cycle (proliferation) and to enter into myogenic differentiation.

We next utilize our dynamic multilayer network analysis to understand the genome reorganization during cellular reprogramming. Here we model time-evolving inter-gene contact maps as a multilayer network, where each layer corresponds to a time instant.

In Fig.7A, we present the overlapping degree (5) of each gene versus its multiplex participation coefficient (6) for networks of four layers at two successive time periods, {0, 8, 16, 24 hrs} and {32, 40, 48, 56 hrs}. Here the overlapping degree provides the total number of contacts associated with a gene, and the multiplex participation coefficient (6) measures the heterogeneity of gene contacts over different layers. We observe that the multiplex participation coefficient has large variance prior to the bifurcation time 32 hrs. In Fig.7B, we extract the top 5% of genes that yield the largest position shift before and after bifurcation in a 2D plane formed by the overlapping degree and the multiplex participation coefficient. As seen from Fig. 7B, most of the genes have smaller multiplex participation coefficients prior to bifurcation, leading to the high degree of heterogeneity of gene contacts over time. In Fig. 7C we show how interactions of genes (e.g., *RPM24, CTGF,* and *FGFBP1*) vary prior to the bifurcation point. We observe that the number of interactions sharply decreases from 8 hrs to 16 hrs, corresponding to a significant change of form prior to bifurcation. This result is consistent with our previous finding shown in Fig.6. Since inter-gene contacts could include both intra-chromosome contacts (the interacting genes belong to the same chromosome) and inter-chromosome contacts (the interacting genes belong to different chromosomes), in Fig. 7D we compare these two types of interactions for genes identified in Fig. 7B. As we can see, the intra-chromosome interaction dominates the genome reorganization during cellular reprogramming.

## 5 Conclusion

We developed network representations of dynamical biological systems based on genome-wide structure and gene expression data captured over time. We also proposed a dynamic multilayer network approach to study the functional organization of the human 4DN. Using graph entropy, we identified critical transitions after which the system shifts abruptly from one state to another, e.g., an intermediate state that is highly receptive and entropic during cellular reprogramming. The use of network centrality helped to identify important topological properties of genome structure and facilitated our integrative form-function analysis based on Hi-C and RNA-seq data. Furthermore, the proposed dynamic multilayer network approach captures the time component that is crucial to advancing our ability to understand cell behavior on a system wide level. In the cellular reprogramming experimental data, we found that nuclear reorganization occurs at the time of cell type specification and that it precedes and facilitates activation of the transcriptional program associated with reprogramming. We believe that there is a general relationship between network entropy and network controllability for a dynamic genome. This would allow us to construct a multi-way dynamical system from genome-wide form-function data. This is a worthwhile topic for future work.

**Fig. 5.**
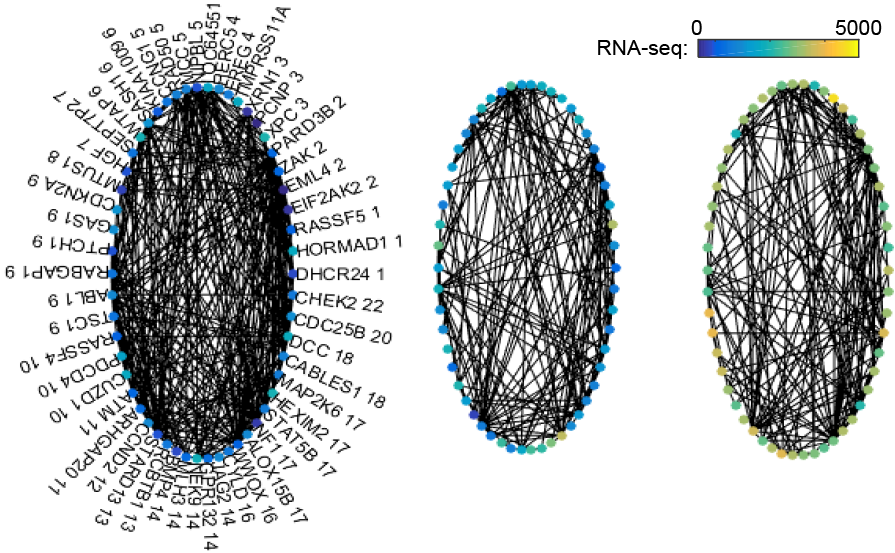
Form-function evolution of selected cell-cycle genes at 0, 16 and 40 hrs (left to right), where each gene marked in the left plot is named as gene name plus chromosome index.

**Fig. 6.**
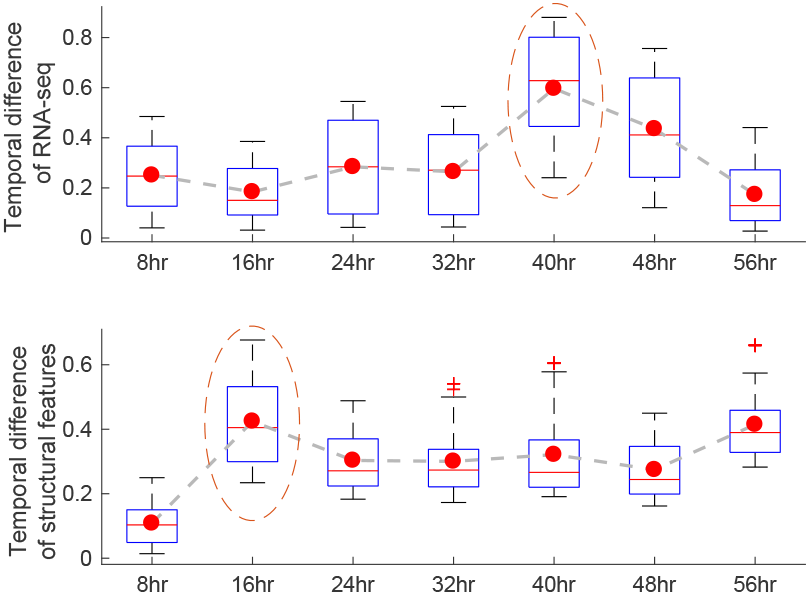
Temporal difference of genomic form and function during reprogramming.

**Fig. 7.**
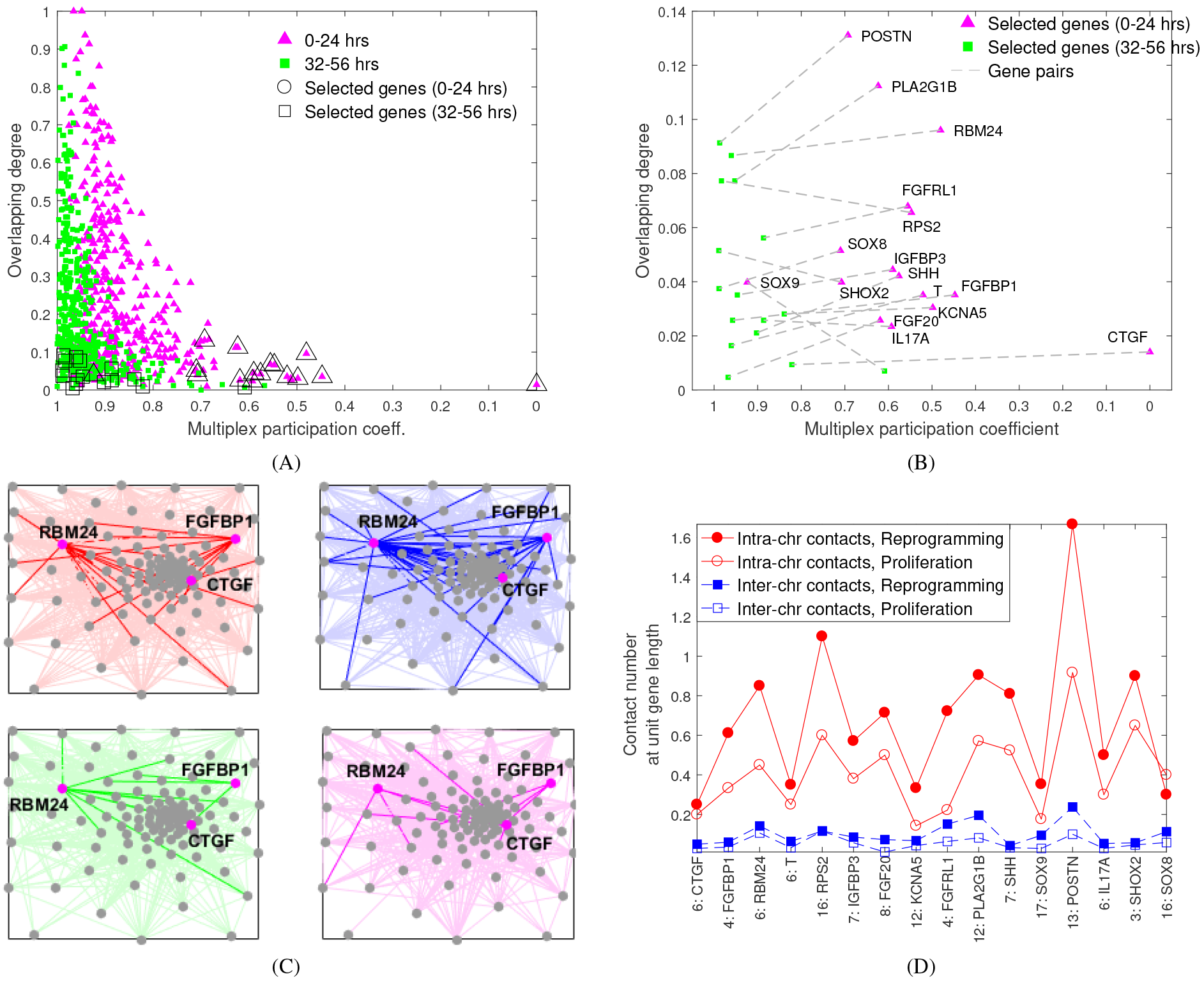
Evolution of inter-gene contact maps during cellular reprogramming. A) Overlapping degree versus multiplex participation coefficient over periods ²0; 8; 16; 24 hrs} and {32; 40; 48; 56 hrs}. B) The top 5‥ of genes yielding the largest position shift before and after bifurcation. c) Time-evolving contacts of representative genes RPM24,CTGF, and FGFBP1 prior to bifurcation. d) Summary of intra-and inter-chromosome contacts of extracted genes.

## Acknowledgements

This work is supported, in part, by the DARPA Biochronicity Program and the DARPA Deep-Purple and FunCC Program. We thank Stephen Lindsly, Haiming Chen and Walter Meixner for their helpful discussions and comments while preparing for the manuscript.

## Multilayer Network Representation

**Fig. S1.**
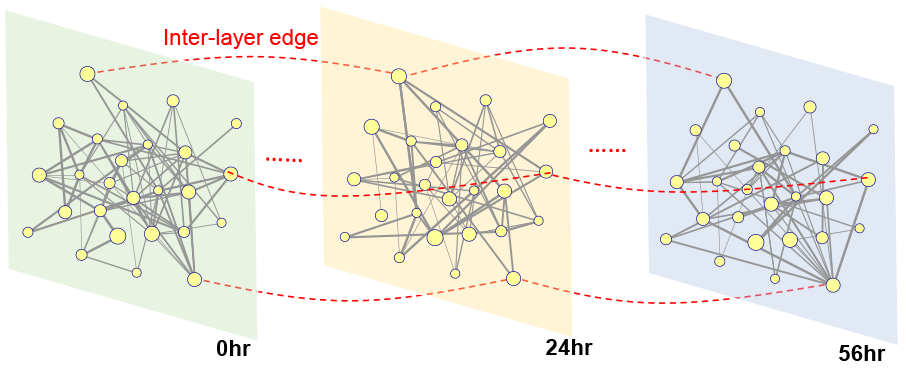
Temporal Hi-C contact maps for 26 cell-cycle genes extracted in (Liu *et al.*, 2017). The inter-layer connection (dash edge) associates one gene to its counterpart in layers before and after the present layer.

### Asymptotic Consistency of VNGE Approximation

#### Theorem

The quadratic approximation Q of VNGE in (2) satisfies

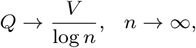

when *n*_+_ ~ *n* and λ_max_ ~ λ_min_, where *n*_+_ denotes the number of positive eigenvalues of L_*c*_, λ_*max*_ and λ_*min*_ denote the largest and smallest nonzero eigenvalues of L_*c*_, and for two functions *f*(*n*) and *g*(*n*) ≠ 0,*f*(*n*) ~ *g*(*n*) means limn_*n*→∞_ *f*(*n*)/*g*(*n*) = 1.

### Proof

Let λ_max_ and λ_min_ denote the largest and smallest nonzero eigenvalue of L_c_, respectively. Assuming L_c_ has at least two nonzero eigenvalues, which implies 0<λ_*i*_ ≤ λ_max_<1 for any nonzero eigenvalue λ_*i*_. Following the definition of *V*, we can rewrite *V* as

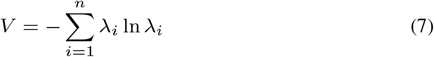

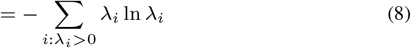

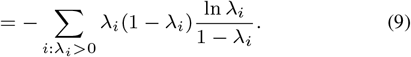

since for all λ_*i*_ > 0, In λ_i_λ_max_ < 0 and 0 < 1-λ_max_≤ 1 - λ_*i*_ ≤ 1 -λ_min_ < 1,we obtain the relation

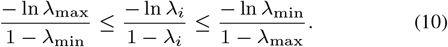

using 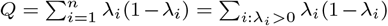and applying to (10) to (9) yeilds

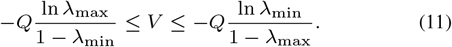

Since 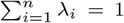 the condition λ_max_ ~λ_min_ implies λ_max_ and λ_min_ are of the same order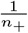, where *n*_+_ is the number of positive eigenvalues of L_*c*_.When the condition *n*_+_ ~*n* also holds, then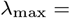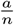 and 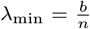 for some constants a,b such that a≥b>0,and we obtain

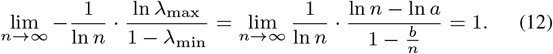

similarly

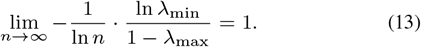

Taking the limit of 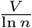and applying (12) and (13) to the bounds in (11),we obtain

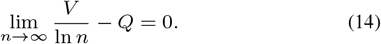

The above Theorem implies that the asymptotic consistency of *Q* with respect to *V* is guaranteed up to a constant factor log *n*. The condition *n*_+_ ~ *n* implies that the number of disconnected components (given by n - n+) is ultimately negligible compared to *n*. The condition λ_max_ ~ λ_min_ implies that a graph Laplacian matrix has balanced eigenspectrum. This condition holds in regular and homogeneous random graphs.

### Dimension Reduction: Laplacian Eigenmap

Laplacian eigenmap is a non-linear dimensionality reduction technique to find a low-dimensional data representation by preserving local properties of the underlying manifold. We remark that the linear dimensionality reduction technique, principal component analysis (PCA), is also applicable but it cannot adequately handle the nonlinearity embedded in the dataset. The method of Laplacian eigenmaps contain the following steps

- Normalize dataset 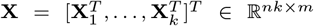 to make different comparable

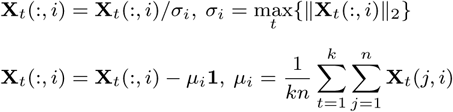

for *i* = 1,2,…,*m*, where *n* is the network size, *k* is the length of time period, *m* is the number of features, **X**_*t*_(:,*i*) denotes the *i*th column of **X**_*t*_, the first transformation ensures that different features are all treated on the same scale, and the second transformation is to zero out the mean of the data.
- Construct a neighborhood graph in which every node is linked with its nearest neighbors. The edge weight is computed using the heat kernel function, leading to a sparse adjacency matrix **W** with entries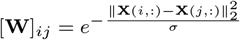, if there is an edge between *i* and *j*, where σis the heat kernel parameter, and we choose σ= 50 in our experiments.
- Compute the graph Laplacian matrix **L** = **D** - **W**, where **D** =diag(BOL
). We then solve the generalized eigenvalue problem Ly = λDy for m` smallest nonzero eigenvalues. The resulting eigenvectors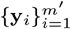form the low-dimensional data representation **Y** = [y_1_, … ym` ] of dimension m`.

### Gene Modules Extracted from Gene Ontology

Genes of interest (GOIs) are mainly extracted through Gene Ontology (GO), with a few GOI subsets curated through other means. GO-extracted lists include muscle, fibroblast, and cell cycle. Here muscle genes are the union of myoblast, myotube, and skeletal muscle genes. The set of genes is given in Supplementary Table 1.

